# Metabolic process of raffinose family oligosacharrides during cold stress and recovery in cucumber leaves

**DOI:** 10.1101/160051

**Authors:** Man Lu, Zhiping Zhang, Jinjin Xu, Wenhua Cao, Minmin Miao

## Abstract

Raffinose family oligosacharrides (RFOs) accumulate under stress conditions in many plants and have been suggested to act as stress protectants. To elucidate the metabolic process of RFOs under cold stress, levels of RFOs and relative carbohydrates, the expression and activities of main metabolic enzymes and their subcellular compartments were investigated during low temperature treatment and recovery period in cucumber leaves. Cold stress induced the accumulation of stachyose in vacuoles, galactinol in vacuoles and cytosols, and sucrose and raffinose in vacuoles, cytosols and chloroplasts. After cold stress removal, levels of these sugars decreased gradually in respective compartments. Among 4 galactinol synthase genes (*CsGS*), *CsGS1* was not affected by the cold stress, while other three *CsGSs* were up-regulated by the low temperature. RNA levels of *acid-α-galactosidase (GAL) 3, alkaline-α-galactosidase (AGA) 2* and *3*, and the activities of GAL and AGA were up-regulated after cold stress removal. The GAL3 protein and GAL activity were exclusively located in the vacuole, whereas the protein of AGA2 and AGA 3 were found in the cytosol and chloroplast respectively. The results indicate that RFOs accumulated during the cold stress in different subcellular compartments in cucumber leaves could be catabolized *in situ* by different galactosidases after stress removal.

## Introduction

Raffinose family oligosaccharides (RFOs) are galactosyl extensions of sucrose that exist widely in the plant kingdom. The physiological functions of RFOs in higher plants have been studied detailedly in the past decades. RFOs are important storage carbohydrates in various plant tissues including leaves, stems, tubers, fruits and seeds, temporarily or terminally (Keller and Pharr, 1996; ElSayed *et al.*, 2014; Sengupta *et al.*, 2015; Ivamoto *et al.*, 2017). In addition, RFOs are used for phloem transport in some plants in Cucurbitaceae, Lamiaceae, Oleaceae, Scrophulariaceae and other several families (Keller and Pharr, 1996; ElSayed *et al.*, 2014; Sengupta *et al.*, 2015).

The important role of RFOs in the stress defence mechanism was also well established. RFOs are characterised as osmoprotectants or antioxidants, and may serve as signals in response to several abiotic or biotic stresses (Zuther *et al.*, 2004; ElSayed *et al.*, 2014; Sengupta *et al.*, 2015). In most sucrose-translocating plants, like rice (*Oryza sativa*) and Arabidopsis (*Arabidopsis thaliana*) that neither transport nor accumulate large quantities of RFOs in their tissues under normal conditions, the accumulation of RFOs and the induced expression of their biosynthetic enzymes, galactinol synthase (GS) and raffinose synthase (RS), were found in response to diverse abiotic stresses such as temperature extremes, drought and salinity (Nishizawa *et al.*, 2008; Saito and Yoshida, 2011; Gangl and Tenhaken, 2016).

Under stressed conditions, the increase of RFOs was also observed in RFOs-translocating plants. *Ajuga reptans*, a frost-hardy perennial labiate, accumulates much more RFOs in winter leaves than in summer leaves, and cold treatment would significantly increase the RFOs concentration in leaves (Bachmann *et al.*, 1994). In cucumber (*Cucumis sativus* L.), *RS* expression, RS activity and the content of raffinose and stachyose increased gradually in the leaves, fruits, stems and roots under low temperature stress (Meng *et al.*, 2008; Sui *et al.*, 2012). In these RFOs-translocating species, it seems that there are two pools of RFOs: a storage pool in the mesophyll (long-term in *Ajuga reptans* or short-term in cucumber), which is involved in stress response, and a transport pool in the phloem (Bachmann and Keller, 1995; Sui *et al.*, 2012).

GS is a key enzyme catalyzing the first step in the RFOs biosynthetic pathway (Keller and Pharr, 1996). Most plants have more than one isoform of GS coded by different genes. In *Ajuga reptans*, there are two GS genes, *ArGolS1* and *ArGolS2. ArGolS1* is mainly involved in the synthesis of storage RFOs while *ArGolS2* is for the synthesis of transport RFOs (Sprenger and Keller, 2000). In the cucumber genome, 4 putative GS genes were found (Wang *et al.*, 2016). However, the exact roles of these genes in the stress response and phloem transport are not well investigated.

The subcellular localization of RFOs and their biosynthetic enzymes under low temperature stress were further studied in *Ajuga reptans* and Arabidopsis, the results showed that GS, RS and stachyose synthase (STS) were extravacuolar (most probably cytosolic), galactosyltransferase, stachyose and higher RFOs were vacuolar, and sucrose and raffinose were found in cytosol, vacuole and chloroplast (Bachmann and Keller, 1995; Tapernoux-Lüthi *et al.*, 2007; Schneider and Keller, 2009; Knaupp *et al.*, 2011; Findling *et al.*, 2015). It is suggested that raffinose, rather than stacyose, plays an important role in stabilizing photosystem II in chloroplasts during low temperature stress in Arabidopsis (Iftime *et al.*, 2011; Knaupp *et al.*, 2011). As a crop of subtropical origin which translocate RFOs but not store large quantities of RFOs under the normal condition, does cucumber have different RFOs subcellular localization with the frost-hardy RFOs-translocating plant *Ajuga reptans* and the sucrose-translocating plant Arabidopsis under cold stress remains unknown.

In contrast to cold acclimation, cold de-acclimation is an important regulatory mechanism to ensure plants restoring to their normal growth state when the stress condition was removed. Unfortunately, although the accumulation of RFOs and its physiological significance under stress conditions have been well studied in several plants, how RFOs were catabolized after stress removal has received little attention. Alpha-galactosidases are responsible for the terminal galactose residue removing during RFOs catabolism (Keller and Pharr, 1996). There are 6 putative α-galactosidase genes in the cucumber genome. These genes are divided into 2 groups, 3 acid α-galactosidase genes (*GAL*) and 3 alkaline α-galactosidase genes (*AGA*), according to their activity in response to pH (Wang *et al.*, 2016). GALs are considered to be localized in the apoplast space or vacuole, while AGAs are supposed to be localized in the cytosol (Keller and Pharr, 1996; Tapernoux-Lüthi *et al.*, 2007). Considering both RFOs and α-galactosidases reveal multiple subcellular localizations, it is interesting to know if different α-galactosidases catabolize the RFOs in different subcellular compartments when stress conditions are relieved.

In this study, in order to reveal the metabolic process of RFOs during cold stress, levels of RFOs and relative carbohydrates, the expression and activities of metabolic enzymes and their subcellular compartments were investigated during cold treatment and the recovery period in cucumber leaves. We emphasized the observation of the expression pattern, activity and intracellular localization of 6 α-galactosidases to elucidate the catabolic process of RFOs after stress removal.

## Materials and methods

### Plant material and temperature treatment

Cucumber (*Cucumis sativus* L.) cultivar Jinchun 5 (from Tianjin Cucumber Institute, China) was used in this study. Seedlings were grown in 10×10 cm plastic pots containing a peat–vermiculite mixture (2:1, v/v) in a growth chamber. The seedlings were thinned to one per pot 10 d after germination. Plants were watered once daily and fertilized weekly with the Hoagland nutrient solution. In the growth chamber, the temperatures were 28°C/22°C (day/night) and the relative humidity 70%. Light was provided by high-pressure mercury lamps (Philip HPLN 400 W) at about 700 μmol m^-2^ s^-1^ for 12 h perday (7:00-19:00). Plants for cold treatment were transferred to another chamber in which the temperature was lowered to 15°C/8°C at the 4-leaf stage. After 3-day chilling treatment, the temperature in the chamber was restored to 28°C/22°C. Control plants remained in the original chamber throughout the experiment. The second leaves from the apical meristem of each plant were collected at 16:00 everyday from the day before treatment to the third day after cold stress removal (named C0, C1, C2, C3, R1, R2, R3, respectively). Samples were frozen in liquid nitrogen immediately after harvest and stored at –80 °C.

### Non-aqueous fractionation of leaves

The procedure was conducted according to Nadwodnik and Lohaus (2008) and Krueger *et al.* (2014) with a few modifications. After removing the middle rib and larger veins, the samples were ground to a fine powder in liquid nitrogen in a precooled mortar and then lyophilized at -25°C. The dry leaf powder was suspended in 20 ml of heptane:tetrachloroethylene mixture (density 1.3 g ml^-1^). Sonication was performed for 2 min, with 6× 10 cycles at 65 % power. The sonicated suspension was filtered through a nylon sieve (40 μm). The sample was centrifuged for 10 min at 3,200 × g and 4 °C and the sediment was resuspended again in the heptane:tetrachloroethylene mixture (density 1.3 g ml^-1^). The suspension was added to an exponential heptane-tetrachlorethylene gradient with a density between 1.27 and 1.50 g ml^-1^. After centrifugation for 60 min at 5,000 × g and 4 °C, six fractions were collected, aliquots of which were taken for the determination of the marker enzymes, RFOs metabolic enzymes and RFOs related sugars. The calculation was carried out by the software BestFit (Krueger *et al.*, 2014).

### Carbohydrate assay and enzyme activity determination

Chloroform methanol extracts were prepared from the aliquots mentioned above for the determination of the carbohydrate concentrations (Nadwodnik and Lohaus, 2008). Galactinol, stachyose, raffinose, galactose, and sucrose were analyzed by HPLC methods as described previously (Miao *et al.*, 2007). For enzymes assay, fractions from gradient centrifugation were washed by 3 volumes of C_7_H_16_ and lyophilized. The dried samples were extracted by Hepes buffer (50 mM Hepes-NaOH pH 7.4; 5 mM MgCl_2_; 1mM EDTA; 1 mM EGTA; 0.1 % Triton X-100; 10% glycerol; 2mM benzamidine; 2mM aminocaproic acid; 1.5mM PMSF; 1g l^-1^ PVPP) (Krueger *et al.*, 2014). Activities of GS and α-galactosidases were assayed according to (Wang *et al.*, 2016). For the assay of RS, the reaction buffer contained 50 mM HEPES–NaOH (pH 7.0), 1 mM DTT, 10 mM galactinol and 40 mM sucrose. Mixtures were incubated at 30 °C for 3 h and the reactions were stopped by boiling for 5 min. The mixture was centrifuged at 28,000×g for 5 min and the supernatant was passed through a 0.45 μm filter. The content of raffinose was determined by HPLC. Enzyme activity is given as μmol of raffinose formation per hour (Sui *et al.*, 2012). The assay of STS was the same as that of RS, except galactinol was replaced by raffinose in the reaction system.

### Total RNA isolation and expression analysis of RS and STS

Total RNA was extracted from approximately 100 mg of the leaf tissues (without middle ribs and larger veins) using TRIzol reagent (Invitrogen, Shanghai, China). Reverse transcription was performed using a Prime Script TM RT reagent Kit with gDNA eraser (Perfect Real Time, TaKaRa, Dalian, China). Quantitative real-time PCR was performed using the One Step SYBR PrimeScript RT-PCR Kit (TaKaRa, Dalian, China) on an ABI PRISM 7700 Sequence Detection System (Applied Biosystems, Shanghai, China), following the manufacturer’s instructions. The real-time PCR was carried out according to the following protocol: 2 min at 94°C, followed by 39 cycles of 94°C for 15 s, 60°C for 15 s and 72°C for 30 s. The cucumber 18S rRNA gene (Gene bank accession No.: AF206894.1) was used for normalization in all the analyses performed. The primer sequences for *RS* and *STS* (Gene bank accession No.: EU096498) are listed in Supplementary Table S1. Primers were confirmed to be approximately 90% to 100% efficient in amplification, and 2^-ΔΔCT^ method (Livak and Schmittgen, 2001) was used for analyses.

### Northern blot analysis of the expression of GS and α-galactosidases genes

Total RNA (20μg) was extracted as mentioned above and electrophoresed on a 1.2% (w/v) agarose gel containing formaldehyde, then transferred onto positively charged nylon membrane (Amersham-Pharmacia Biotech, Uppsala, Sweden) with 10×SSC buffer (1×SSC containing 150 mM NaCl and 15 mM sodium citrate, pH 7.0). DNA fractions of 4 Cs*GS* genes and 6 α-galactosidases genes were cloned into pMD 18-T vector (Takara, China) as DNA templates for RNA probe synthesis. The primer sequences for cloning and the gene bank accession No. of 10 genes are listed in Supplementary Table S1. The probe synthesis and northern blot was performed using DIG Northern Starter Kit (Roche Applied Science, China) following the manufacturer’s instructions.

### Subcellular localization of 6 α-galactosidases

To obtain the “cauliflower mosaic virus (CaMV) 35S promoter::GFP::gene ORF” fusion protein expression structure, the CaMV 35S promoter sequence was amplified with primers 5’- ccggaattccatggagtcaaagat -3’ (*EcoR* I restriction site added) and 5’- ataaggatccagtcccccgtgtt -3’ (*BamH* I restriction site added) from the vector pCambia1303. The enhanced GFP (EGFP) sequence was amplified with primers 5’-aatgtcgacatggtgagcaagggcgagg-3’(Sal I restriction site added) and 5’-gaatctgcagcttgtacagctcgtccatg-3’ (*Pst* I restriction site added) from the vector pEGFP-N1(U55762). These two fragments were inserted into the vector pCambia1381*c using restriction enzymes mentioned above. All α-galactosidase genes were further cloned into the vector “pCambia1381*c:: CaMV 35S:: EGFP”. The primers and restriction enzymes used are list in the Supplementary Table S2. The construction of *GAL3* fusion protein expression system is the same as other 5 α-galactosidase genes except the restriction enzymes *Apa* I and *EcoR* I were used for CaMV 35S promoter cloning, and *EcoR* I and *BamH* were used to clone EGFP fragments.

Transient transformation of onion epidermis was carried out according to Xu *et al.* (2004) with modifications. The vectors pCambia1381*c containing “CaMV 35S promoter::EGFP::gene ORF” structures were transformed into *Agrobacterium tumefaciens* strain LBA4404. *Agrobacterium* cultivated overnight at 28°C was harvested at OD_600_ of 1.5 to 2.0, centrifuged at 5000 rpm for 10 min and resuspended in 50 ml of infiltration liquid (50g l^-1^ sucrose+100mg l^-1^ acetosyringone). Adaxial onion epidermis 1 cm^2^ was incubated in the infiltration liquid for 30s and then cultured on the 1/2 MS solid medium (30g l^-1^ sucrose, 7g l^-1^ agar, pH 5.8) for 2 d ( 25°C, 16h/8h). Plasmolysis cells were obtained by adding 300g l^-1^ KNO_3_ on the tissue for 5min before fluorescence microscope observation (Zeiss LSM710). Transient transformation of tobacco leaves was mainly according to Sparkes *et al.* (2006). Basically, the underside of the tobacco leaf was punctured with a small needle. Took up resuspended *Agrobacterium* infiltration liquid (10 mM MgCl_2_, 10 mM MES, 100μM acetosyringone) in a 1 ml syringe (no needle). Placed the tip of the syringe against the underside of the leaf over the needle mark and pressed down gently on the plunger. Placed plants in a growth cabinet under normal conditions for 3d, and then excised 1-2 cm^2^ segments of leaf tissue for confocal microscope observation (Zeiss LSM710).

## Results

### Subcellular distribution of sugars and relative enzymes in cold-stressed cucumber leaves

Since RFOs will accumulate after cold stress (Sui *et al.*, 2012), cucumber leaves were sampled after a 3-day cold treatment. The distribution of the sugars and relative enzymes among the vacuolar, chloroplast, and cytosolic compartment was measured by non-aqueous fractionation method and the results are summarized in the Table 1. Stachyose and galactose were almost exclusively found in the vacuole, sucrose and raffinose were distributed in all three compartments, while galactinol was located mainly in the vacuole and cytosol. Three RFOs synthesis enzymes, GS, RS and STS, were mostly located in the cytosol. The GAL activity was mainly found in the vacuole whereas the AGA activity was mostly distributed in both chloroplast and cytosol (Table 1).

**Table 1.** Percentage distribution of carbohydrates and enzymes of RFOs metabolism among the cytosols, chloroplasts and vacuoles of leaf mesophyll cells in cucumber seedlings under cold stress for 3 days. The data represent the mean ±SE of three samples.

| Carbohydrates/enzymes | Cytosol (%) | Chloroplast (%) | Vacuole (%) |
| --- | --- | --- | --- |
| Galactinol | 53.2±6.3 | 9.4±4.2 | 38.4±10.2 |
| Sucrose | 37.2±17.2 | 40.2±13.8 | 22.6±8.4 |
| Raffinose | 42.1±12.4 | 21.8±9.6 | 36.1±15.3 |
| Stachyose | 1.7±1.3 | 2.1±1.7 | 96.2±3.6 |
| Galactose | 0.9±1.4 | 0.8±2.3 | 98.3±4.3 |
| Acid-α-Galactosidase | 3.2±0.8 | 1.3±0.5 | 95.5±7.2 |
| Alkaline-α-Galactosidase | 66.3±8.3 | 27.4±5.6 | 6.3±3.2 |
| Galactinol synthase | 100±0.0 | 0.0±0.0 | 0.0±0.0 |
| Raffinose synthase | 100±0.0 | 0.0±0.0 | 0.0±0.0 |
| Stachyose synthase | 99.1±2.6 | 0.0±0.0 | 0.9±1.2 |

***Suggested position of Table1***

### Change of sugar levels during cold treatment and recovery

According to the subcellular distribution of sugars (Table 1), levels of galactinol, sucrose, raffinose, stachyose and galactose in vacuole, sucrose and raffinose in chloroplast, and galactinol, sucrose and raffinose in cytosol were investigated during the cold stress and recovery period (Fig. 1). The levels of all measured sugars except galactose increased from 0 d to 3 d of cold treatment and then decreased when the temperature was recovered to the normal level, indicating low temperature promoted the accumulation of these sugars in different subcellular compartments. The galactose level showed an opposite pattern, which decreased during cold stress and increased during the recovery period in the vacuole. The amplitude of fluctuation of sugar levels was more significant in vacuole and chloroplast than in cytosol. No significant fluctuations of sugar levels were found in control plant leaves during the treatment (Supplementary Fig. S1).

**Fig.1.**
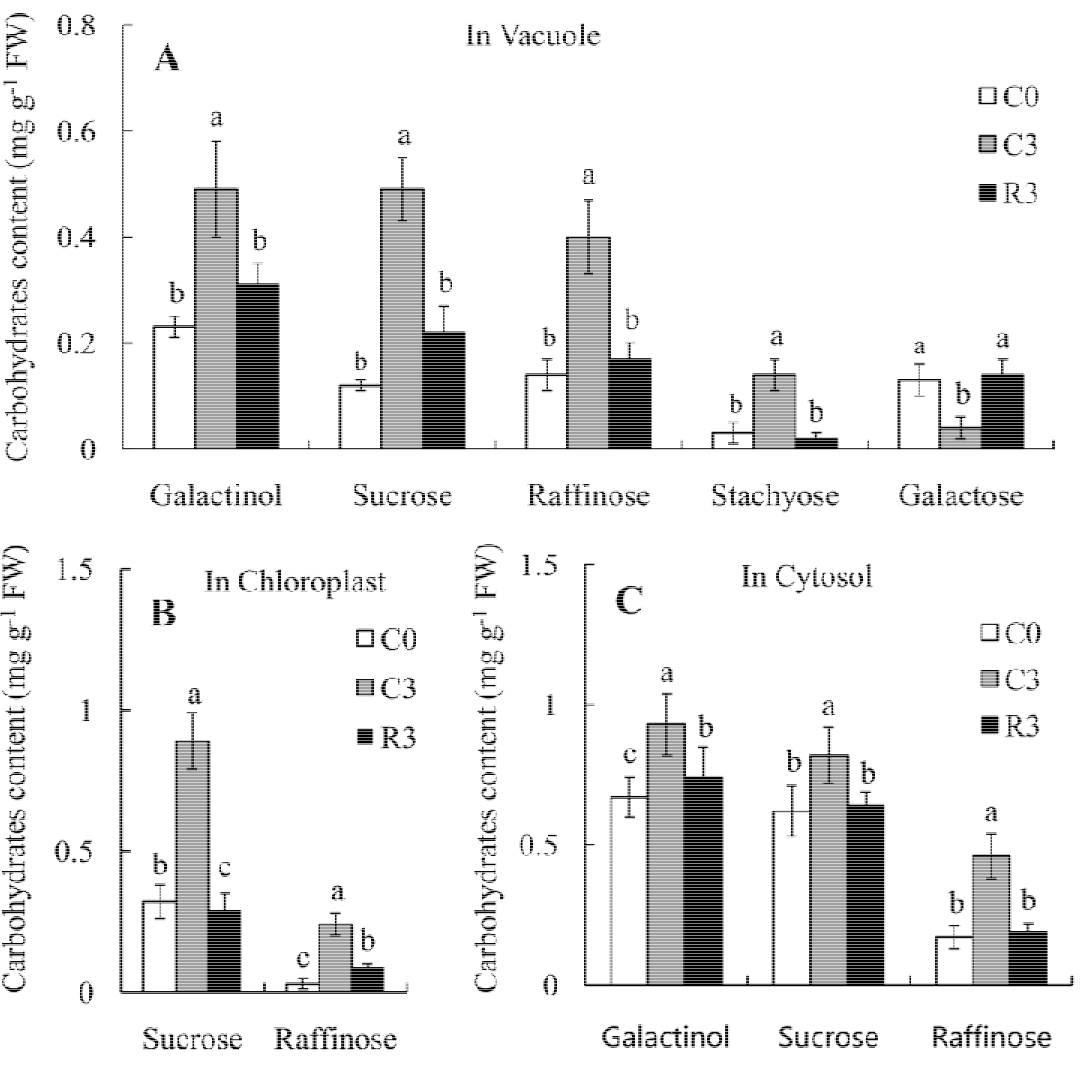
Changes of sugar levels in different subcellular compartments during cold stress and recovery. C0: cold stress for 0 d; C3: cold stress for 3d; R3: the 3rd day after stress removal. Means ±SE (5 samples) followed by different letters are significantly different (P <0.05).

### The expression and activities of RFOs synthesis enzymes during cold treatment and recovery

Four *GSs* showed different expression patterns under cold stress. Under normal temperature, the expressions of *CsGS1* and *CsGS4* were detected and the expression of *CsGS4* was stronger than that of *CsGS1*, while the expressions of *CsGS2* and *CsGS3* were not detected by the northern blot. Cold treatment had no significant effect on the expression of *CsGS1.* However, *CsGS2, CsGS3* and *CsGS4* expressions were enhanced by the low temperature, and RNAs of *CsGS2* and *CsGS3* could be detected at the 3rd day of cold treatment. After the stress removal, the RNAs levels of these three genes declined gradually. At the 3rd day of temperature recovery, the expression of *CsGS3* fell below the detection limit (Fig. 2A). The GS activity was closely correlated to the expression pattern of the *CsGS2, CsGS3* and *CsGS4*, which increased under the cold stress and decreased after the stress removal (Fig. 2B).

**Fig.2.**
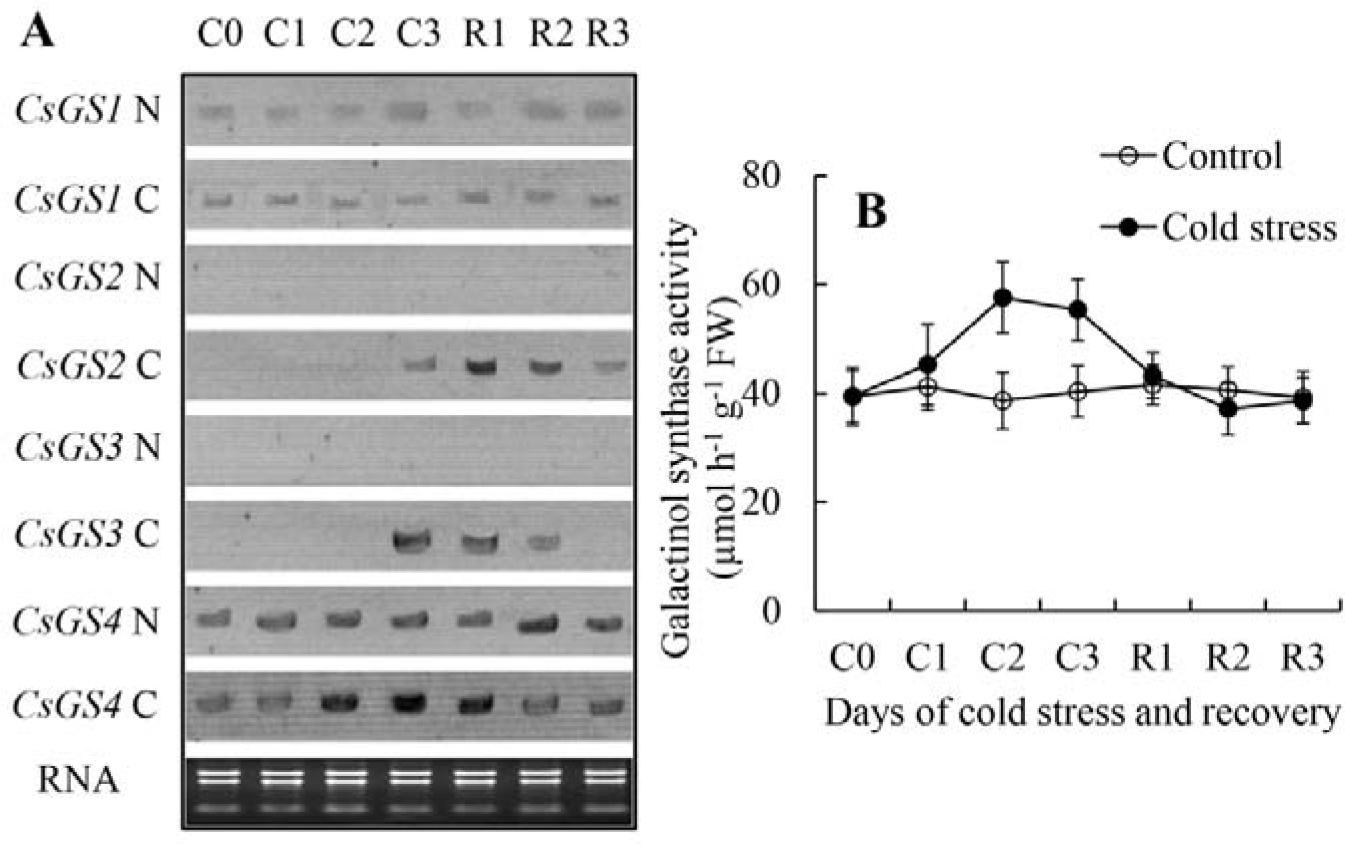
Changes of expression and activity of galactinol synthases (GSs) in cucumber leaves during cold stress and recovery. A: expression; B: activity. N: normal temperature (28°C/22°C); C: Cold stress (15°C /8°C); C0, C1, C2, C3: cold stress for 0 d, 1 d, 2 d and 3 d; R1, R2, R3: temperature was restored to normal level for 1 d, 2 d and 3 d. Each point is the average of 5 samples. Error bars represent SEs.

The *RS* showed a similar expression pattern to that of the *CsGS2, CsGS3* and *CsGS4.* The RNA level of *RS* was significantly up-regulated by the low temperature, and then declined after the stress removal (Fig. 3A). The RS activity also increased, and was higher than that of control plant at the 3rd day of recovery (Fig. 3B). No clear effect of cold stress on the STS expression and enzyme activity was observed in this study (Fig. 3C, D).

**Fig.3.**
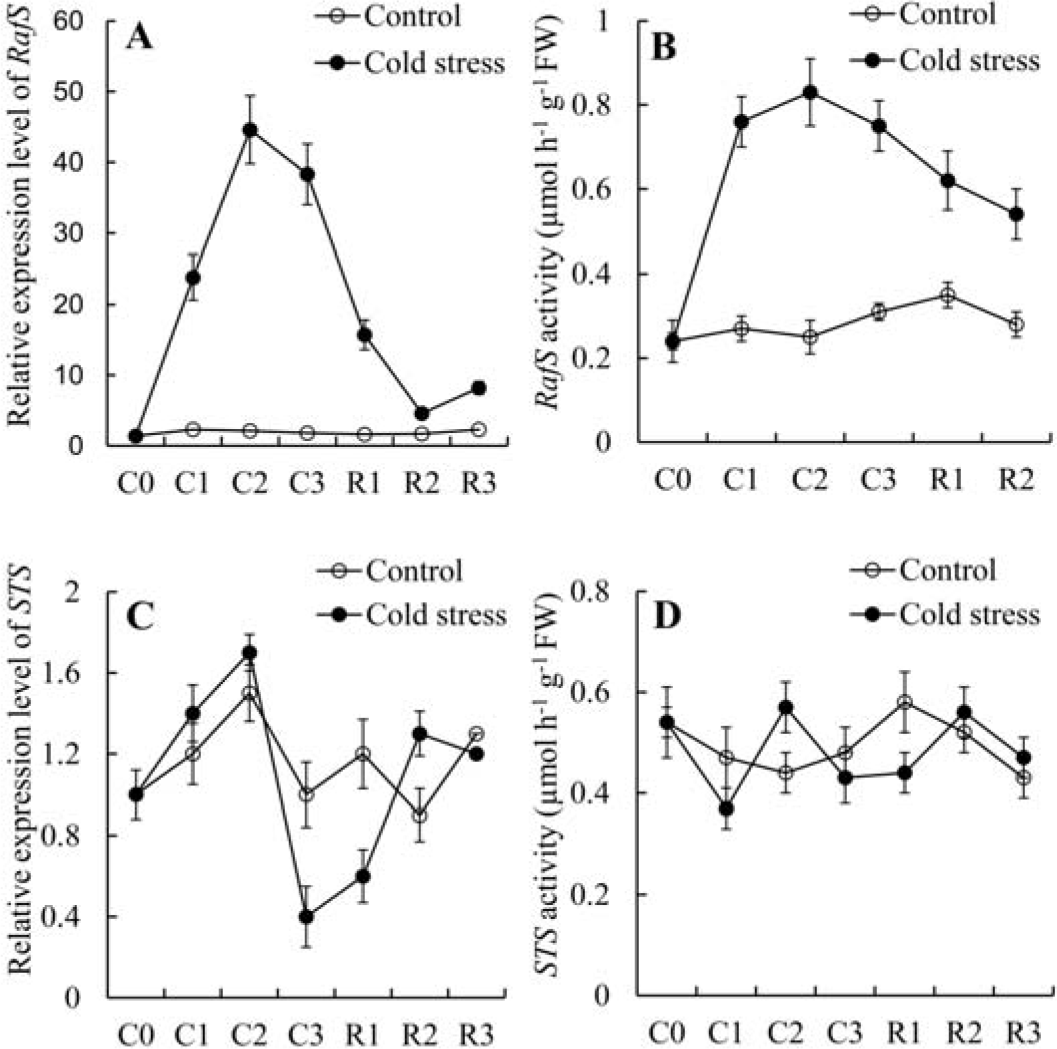
Changes of expression and activity of raffinose synthase (RS) and stachyose synthase (STS) in cucumber leaves during cold stress and recovery. A: RS expression; B: RS activity; C: STS expression; D: STS activity. Control: 28°C/22°C; Cold stress: 15°C/8°C; C0, C1, C2, C3: cold stress for 0 d, 1 d, 2 d and 3 d; R1, R2, R3: temperature was restored to normal level for 1 d, 2 d and 3 d. Each point is the average of 5 samples. Error bars represent SEs.

### The expressions and activities of galactosidases during cold treatment and recovery

As shown in Fig. 4A, among 3 *GALs*, only *GAL1* RNA was detected under normal temperature, and low temperature treatment and recovery have no significant effect on its level. *GAL2* mRNA was not detected under both treatments. The expression of *GAL3* was undetectable under both normal and low temperature, but was induced remarkably when the cold stress was removed. mRNAs of all 3 *AGAs* were detected under the normal temperature. No significant effect of temperature change on the *AGA1* expression was found. However, the RNA levels of both *AGA2* and *AGA3* were down-regulated by the cold stress and then restored to normal levels after cold stress removal (Fig. 4 B). Both GAL and AGA activities were measured in different subcellular compartments (Fig. 5). Two substrate, raffinose and stachyose were used in the enzyme assay. The GAL activity was higher with the substrate raffinose, while AGA activity was higher with stachyose. The fluctuation patterns of enzyme activities with two substrates were similar in all treatments. In vacuole, the GAL activity remained unchanged during cold stress and increased significantly after stress removal. In chloroplast and cytosol (Fig. 5A, C), the AGA activities were down-regulated by the low temperature and then up-regulated by the stress removing (Fig. 5 B, D). No significant fluctuation of GAL or AGA activities was found in control plant leaves during the treatment (Supplementary Fig. S2). These results indicated that *GAL3, AGA2* and *AGA3* were important for RFOs catabolism during the temperature recovery.

**Fig.4.**
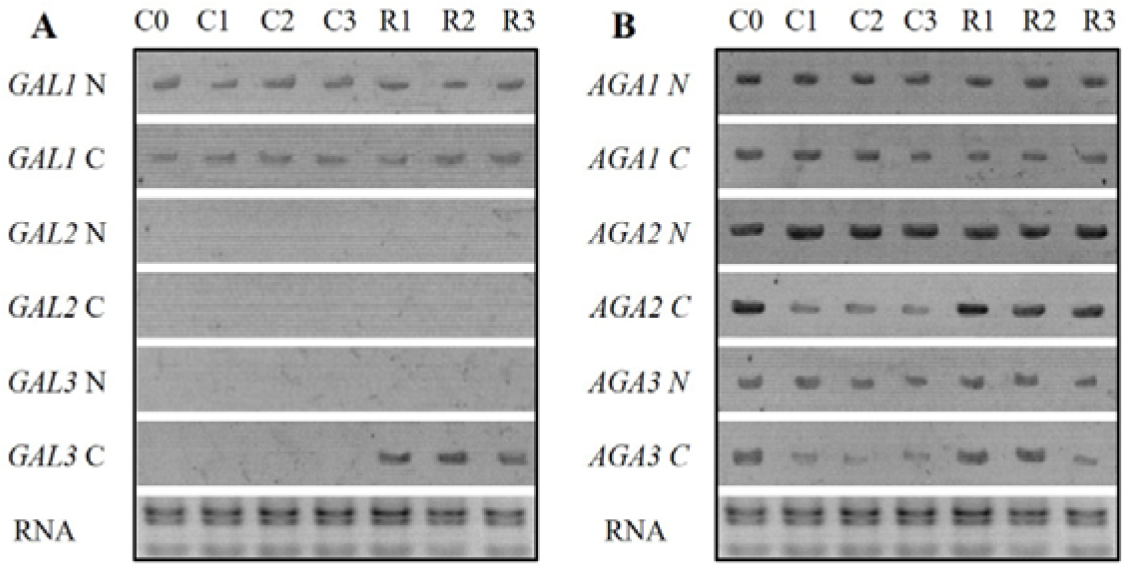
Expression of α-galactosidases in cucumber leaves during cold stress and recovery. A: acid-α-galactosidases (*GAL*); B: alkaline-α-galactosidases (*AGA*); N: Normal temperature (28°C/22°C); C: Cold stress(15°C /8°C); C0, C1, C2, C3: cold stress for 0 d, 1 d, 2 d and 3 d; R1, R2, R3: temperature was restored to normal level for 1 d, 2 d and 3 d.

**Fig.5.**
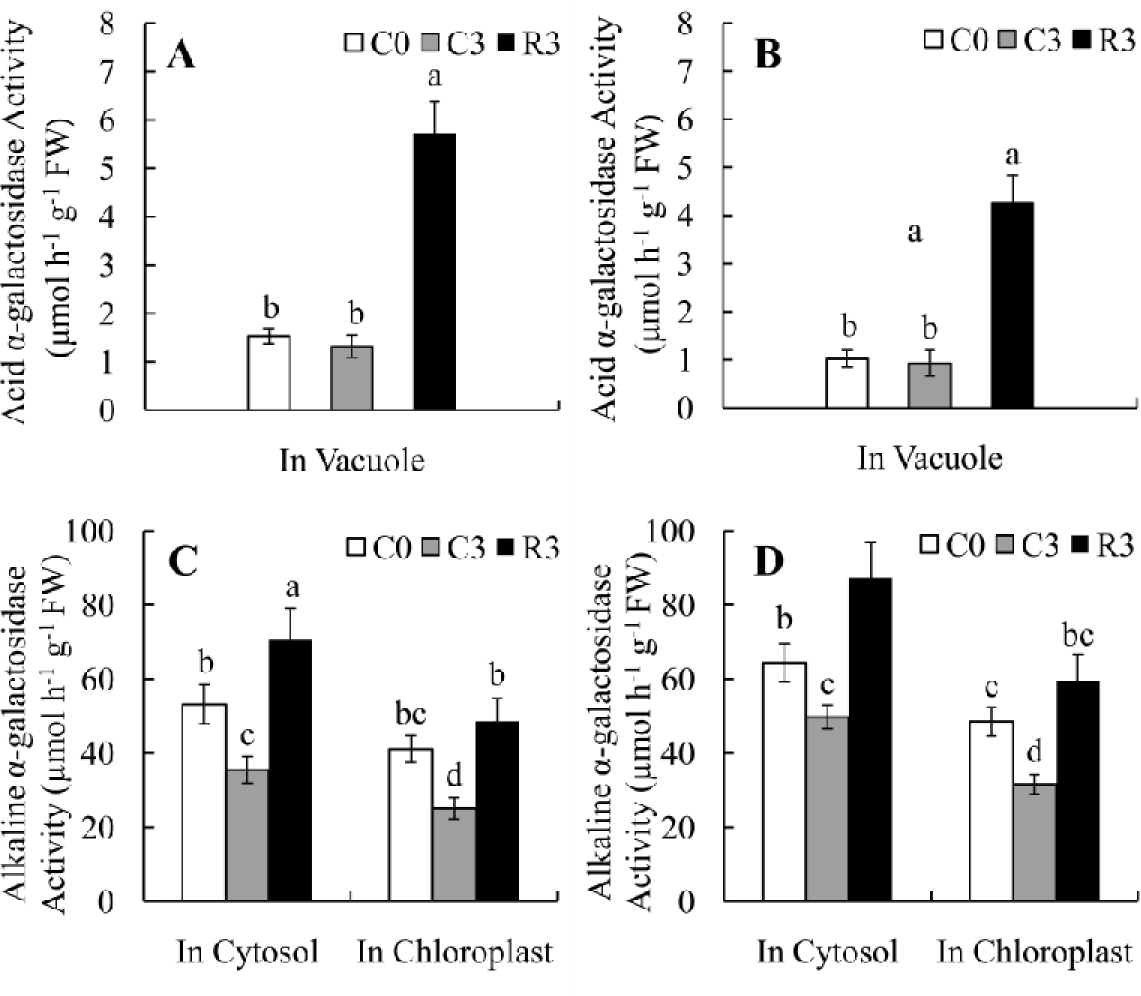
Changes of α-galactosidase activities in different subcellular compartments of cucumber leaves during cold stress and recovery. A and C: using raffinose as substrate; B and D: using stachyose as substrate; C0: Cold stress for 0 d; C3: cold stress for 3d; R3: the 3rd day after stress removal. Means±SE (5 samples) followed by different letters are significantly different (P <0.05).

### Subcellular localization of galactosidase proteins

To obtain further insight into the mechanism of RFOs catabolism after cold stress removal, an EGFP protein was fused to the N-terminus of 6 galactosidases, and placed under the control of the CaMV 35S promoter. These constructs were transiently expressed in the onion epidermal cells. To distinguish the cell wall and the cytosol in these highly vacuolate cells, plasmolysis was carried out before transformation. Fluorescence microscope imaging showed that GAL1 and GAL2 were located near the cell wall, GAL3 was found in the vacuole, while all 3 AGAs were distributed in the cytosol (Fig. 6). The subcellular localization of AGA2 and AGA3, which were found to play a role in catabolizing RFOs in cytosol or chloroplast, were further determined by transiently expressing the EGFP fusion construct in the tobacco mesophyll cell. The results revealed that AGA2 was located in the cytosol, while AGA3 was mostly found in the chloroplast (Fig. 7).

**Fig.6.**
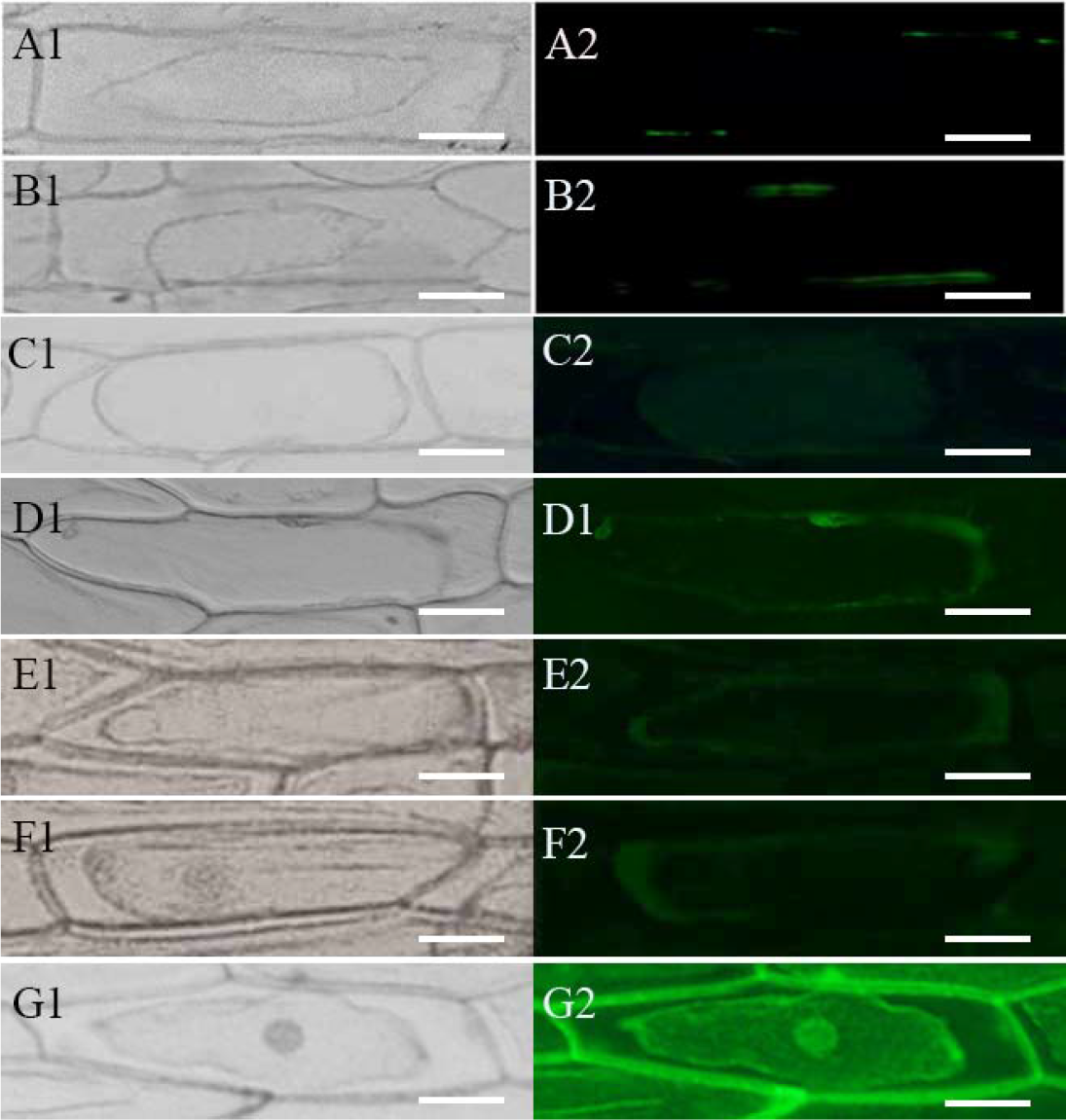
Subcellular localization of acid-α-galactosidases (GALs) and alkaline-α-galactosidases (AGAs). GFP-GAL and GFP-AGA fusion proteins and GFP alone expressed under the control of the CaMV35S promoter in onion epidermal cells were observed under a fluorescent microscope. A1and A2: GAL1; B1and B2: GAL2; C1and C2: GAL3; D1and D2: AGA1; E1and E2: AGA2; F1and F2: AGA3; G1and G2: GFP alone. A1, B1, C1, D1, E1, F1, G1: Differential interference contrast images; A2, B2, C2, D2, E2, F2, G2: GFP fluorescence signals. To distinguish the cell wall and the cytosol in these highly vacuolate cells, plasmolysis was carried out before transformation. Bars=50μm.

**Fig.7.**
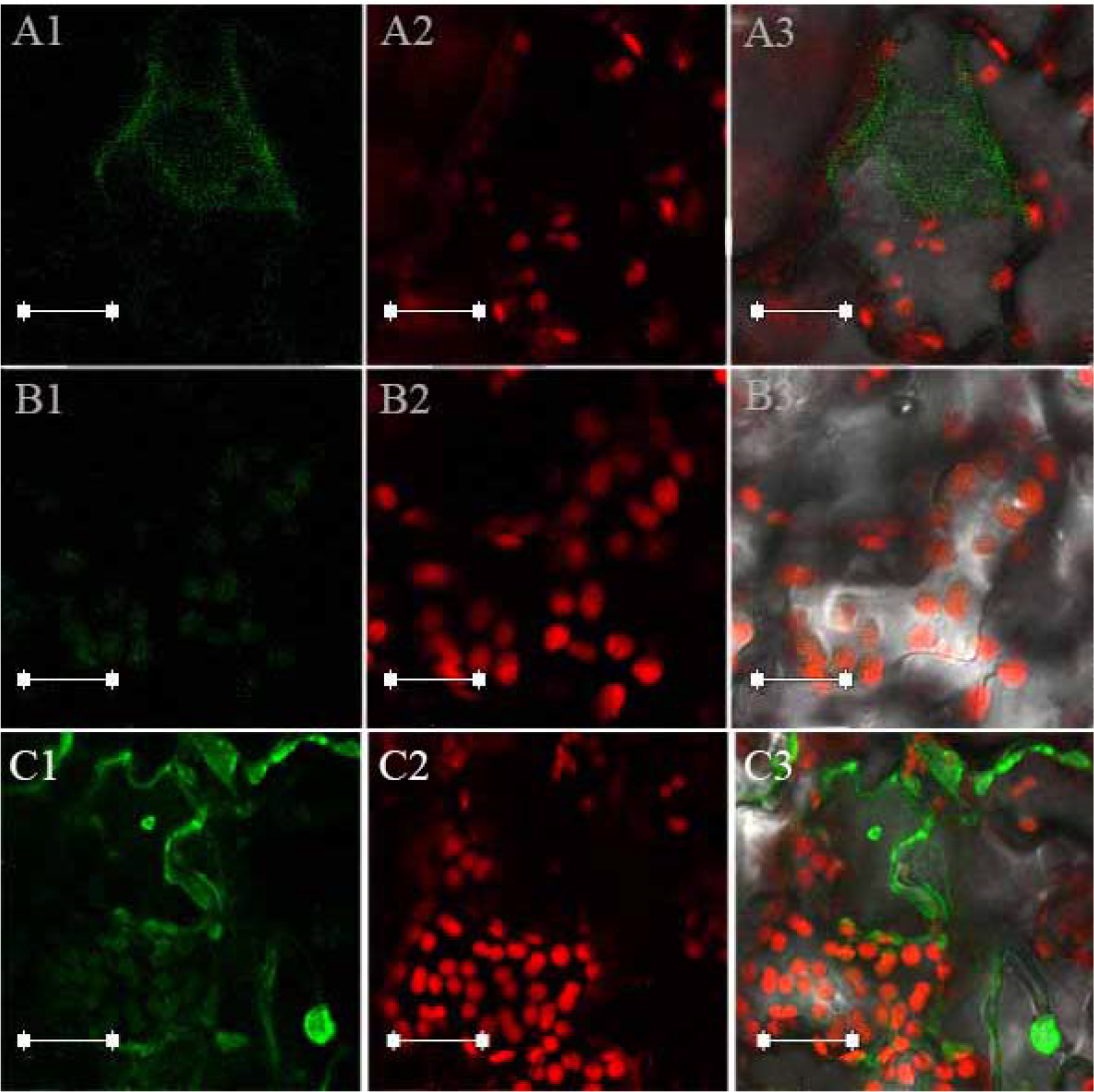
Subcellular localization of alkaline-α-galactosidases (*AGAs).* GFP-AGA2 fusion protein (A1, A2 and A3), GFP-AGA3 fusion protein (B1, B2 and B3) and GFP alone (C1, C2 and C) expressed under the control of CaMV35S promoter in tobacco leaf cells were observed under a confocal microscope. The left column (A1, B1 and C1) shows green channel (GFP signal, 488 nm), the middle column (A2, B2 and C2) shows red channel (chloroplast autofluorescence signal, 633nm), and the right column (A3, B3 and C3) shows merged images. Bars=10μm.

## Discussion

The subcellular compartment study of RFOs and relative carbohydrates after cold stress in cucumber leaves using non-aqueous fractionation method showed similar results with previous reports, *i.e.*, stachyose and galactose were mostly in the vacuole, galactinol in the vacuole and cytosol, and sucrose and raffinose in the vacuole, cytosol and chloroplast (Schneider and Keller, 2009; Knaupp et al., 2011; Nägele and Heyer, 2013; Findling *et al.*, 2015). In cucumber leaves, besides accumulating after cold stress, RFOs are synthesized in the cytosol of intermediary cells in the minor veins for phloem transport (Turgeon and Wolf, 2009). The effect of this “transport RFOs pool” on the results in this study seemed negligible, since little stachyose are found in the cytosol (Table 1). The levels of galactinol, sucrose and raffinose increased under the low temperature treatment and decreased after the stress removal, similar phenomena were also found in other plant species (Cunningham *et al*, 2003; Brenac *et al.*, 2013; ElSayed *et al.*, 2014). The results of this study, together with data from *Ajuga reptans* (Bachmann *et al.*, 1994), suggest that accumulations of RFOs are also important for RFOs translocating species to deal with cold stress. Iftime *et al.* (2011) have shown that stachyose accumulation in transgenic Arabidopsis plants did not increase the freezing tolerance. However, the up-regulated level of this tetrasccharide in the vacuole in cold acclimated cucumber and *Ajuga reptans* leave tissues (Findling *et al.*, 2015; this study) suggest that stachyose may exert its protective role in these RFOs-translocating plants. Unlike *Ajuga reptans*, cucumber does not accumulated higher RFO oligomers in leaves (Meng *et al.*, 2008). The physiological importance of stachyose accumulated in the vacuole during cold stress in cucumber leaves awaits further research.

GSs are always encoded by multiple genes in plant genomes. Evidences indicated that these isoforms have different cellular locations and physiological functions. In Arabidopsis, *AtGolS1* and *AtGolS 2* were induced by drought and high-salinity stresses, while *AtGolS3* was induced by cold stress (Taji *et al.*, 2002). In the stachyose-translocating plant *Ajuga reptans, ArGolS1* expressed in the mesophyll for storage RFOs synthesis and *ArGolS2* mainly in the intermediary cell for transport RFOs synthesis (Sprenger and Keller, 2000). In the melon (*Cucumis melo*), Cucumber mosaic virus and heat stress did not affect the expression level of *CmGolS1*, but caused a significant increase in the relative expression level of *CmGolS2* (Gil *et al.*, 2012). In this study, in cucumber leaves *CsGS1* expressed constitutively and was not affected by the cold stress, while the expressions of other three *CsGSs* were up-regulated by the low temperature. Phylogenetic analysis based on the amino acid sequences indicated that cucumber *CsGS1* and *CmGloS1* are closely related, while cucumber *CsGS4* and *CmGloS2* cluster into one group (Supplementary Fig. S3). It is not clear if the function of *CsGS1* and *CmGloS1* is similar to that of the *Ajuga reptans ArGolS2* (for transport RFOs synthesis). *CsGS2* and *CsGS3* showed similar expression pattern during the temperature treatment, *i.e.*, could not be detected under normal temperature and induced by cold stress. In the phylogenetic tree, cucumber *CsGS2* is closely related to the *SmGloS3*, which has a low constitutive expression and could be induced by several abiotic stresses in *Salvia miltiorrhiza* (Wang *et al.*, 2012). *CsGS3, AtGolS2* and *AtGolS3* belong to the same group (Supplementary Fig. S3). It seemed that the function of *GSs* in abiotic stress is mainly determined by which element exists in their promoter area, rather than the amino acid sequences (Taji *et al.*, 2002). The response patterns of 4 *CsGSs* to other abiotic stresses need further investigation. Other two RFOs biosynthetic enzymes, RS and STS, revealed different expression and activity patterns during temperature treatment, RS was up-regulated by cold stress but STS not. The increased level of substrate raffinose, rather than the biosynthetic enzymes STS, may result in the stachyose accumulation under low temperature. In addition, the cytosol compartment of GS, RS and STS, the chloroplast and vacuole localization of raffinose and the vacuole localization of stachyose, further confirm that there are RFOs transporters on the tonoplast and chloroplast envelope in RFOs-translocaing plants (Greutert and Keller, 1993; Schneider and Keller, 2009; Nägele and Heyer, 2013).

Up to date, little research has been focused on the catabolism process of RFOs after stress conditions are removed in leaf tissues. Subcellular localizations of 6 cucumber galactosidases, which are considered key enzymes in the pathway, were studied in this research. Tapernoux-Lüthi *et al.* (2007) concluded that a C-terminal oligopeptide extension is a non-sequence-specific vacuolar sorting determinant of plant galactan:galactan galactosyltransferase and acid galactosidase. Sequence analysis revealed that among 3 cucumber *GALs*, only *GAL3* has this C-terminal oligopeptide extension (Supplementary Fig. S4), indicating the vacuolar compartment of *GAL3*, and the apoplastic location of *GAL1* and *GAL2.* The results of our EGFP fusion protein transiently expression experiments confirmed that *GAL3* is the only acid galactosidase located in the vacuole. Combining with the expression and activity pattern during the temperature treatment, we concluded that *GAL3* was responsible for the RFOs catabolism in vacuoles after stress removal. Alkaline galactosidases were always considered to be distributed in the cytosol (Keller and Pharr, 1996). *Osh69*, a rice alkaline galactosidase, was found to be located in the chloroplast and play a role during leaf senescence (Lee *et al.*, 2004). In this study, cucumber *AGA3* protein was found in the chloroplast and its expression was down-regulated by the cold stress and up-regulated by the temperature recovery, strongly suggests an important role for *AGA3* in chloroplast RFOs catabolism after cold stress removal. The alkaline environment of chloroplast stroma is suitable for the AGA3 to exert its catalytic function (Findling *et al.*, 2015). The data of this study also indicated that another alkaline galactosidase, AGA 2, is responsible for RFOs catabolism in the cytosol after temperature was recovered to normal level.

In conclusion, our results indicate that RFOs accumulated during cold stress in different subcellular compartments in cucumber leaves could be catabolized *in situ* by different galactosidases after stress removed. The data do not rule out the possibility that RFOs are translocated to other subcellular compartments for degradation.

## Supplementary data

**Fig. S1.** Changes of sugar levels in different subcellular compartments of control plant leaves during treatment.

**Fig. S2.** Changes of α-galactosidase activities in different subcellular compartments of control plant leaves during treatment.

**Fig. S3.** Phylogenetic tree representing the relationship of galactinol synthase genes from different plant species with the full-length protein sequence reported.

**Fig. S4.** Sequence comparison of C-terminal peptides of three cucumber acid alpha-galactosisase genes.

**Table S1.** Primers used for real-time PCR and Northern hybridize probe synthesis.

**Table S2.** Primers used for EGFP fusion protein vector construction.

## Acknowledgements

This work was supported by the National Natural Science Foundation of China (grant no. 31672160).

## References

Bachmann M, Keller F. 1995. Metabolism of the raffinose family oligosaccharides in leaves of Ajuga reptans L. Inter- and intracellular compartmentation. Plant Physiology 109, 991–998.

Bachmann M, Philippe M, Keller F. 1994. Metabolism of the raffinose family oligosaccharides in leaves of Ajuga reptans L. Cold acclimation, translocations, and sink to source transition: discovery of chain elongation enzyme. Plant Physiology 105, 1335–1345.

Brenac P, Horbowicz M, Smith ME, Obendorf RL. 2013. Raffinose and stachyose accumulate in hypocotyls during drying of common buckwheat seedlings. Crop Science 53, 1615–1625.

Cunningham SM, Nadeau P, Castonguay Y, Laberge S, Volenec JJ. 2003. Raffinose and stachyose accumulation, galactinol synthase expression, and winter injury of contrasting alfalfa germplasms. Crop Science 43, 562–570.

ElSayed AI, Rafudeen MS, Golldack D. 2014. Physiological aspects of raffinose family oligosaccharides in plants: protection against abiotic stress. Plant Biology 16, 1–8.

Findling S, Zanger K, Krueger S, Lohaus G. 2015. Subcellular distribution of raffinose oligosaccharides and other metabolites in summer and winter leaves of *Ajuga reptans* (Lamiaceae). Planta 241, 229–241.

Gangl R, Tenhaken R. 2016. Raffinosr family oligosaccharides act as galactose stores in seeds and are required for rapid germination of Arabidopsis in the dark. Frontiers in Plant Science 7, 1115.

Gil L, Ben-Ari J, Turgeon R, Wolf S. 2012. Effect of CMV infection and high temperature on the enzymes involved in raffinose family oligosaccharide biosynthesis in melon plants. Journal of Plant Physiology 169, 965–970.

Greutert H, Keller F. 1993. Further evidence for stachyose and sucrose/H+ antiporters on the tonoplast of Japanese artichoke (Stachys sieboldii) tubers. Plant Physiology 101, 1317–1322.

Iftimea D, Hannah MA, Peterbauer T, Heyer AG. 2011. Stachyose in the cytosol does not influence freezing tolerance of transgenic Arabidopsis expressing stachyose synthase from adzuki bean. Plant Science 180, 24–30.

Ivamoto ST, Reis O Júnior, Domingues DS, dos Santos TB, de Oliveira FF, Pot D, et al. 2017. Transcriptome analysis of leaves, flowers and fruits perisperm of Coffea arabica L. reveals the differential expression of genes involved in raffinose biosynthesis. PLoS ONE 12, e0169595.

Keller F, Pharr DM. 1996. Metabolism of carbohydrates in sinks and sources: galactosyle-sucrose oligosaccharides. In: Zamsk E, Schaffer,AA, eds. Photoassimilate Distribution in Plants and Crops. New York: Marcel Dekker, 157–183.

Knaupp M, Mishra KB, Nedbal L, Heyer AG. 2011. Evidence for a role of raffinose in stabilizing photosystem II during freeze-thaw cycles. Planta 234, 477–486.

Krueger S, Steinhauser D, Lisec J, Giavalisco P. 2014. Analysis of subcellular metabolite distributions within Arabidopsis thaliana leaf tissue: a primer for subcellular metabolomics. Methods in Molecular Biology 1062, 575–596.

Lee RH, Lin MC, Chen SC. 2004. A novel alkaline-galactosidase gene is invovled in rice leaf senescence. Plant Molecular Biology 55, 281–295.

Livak KJ, Schmittgen TD. 2001. Analysis of relative gene expression data using real-time quantitative PCR and the 2^-ΔΔCT^ method. Methods 25, 402–408.

Meng FZ, Hu LP, Wang SH, Sui XL, Wei L, Wei YX, Sun JL, Zhang ZX. 2008. Effects of exogenous abscisic acid (ABA) on cucumber seedling leaf carbohydrate metabolism under low temperature. Plant Growth Regulation 56, 233–244.

Nägele T, Heyer AG. 2013. Approximating subcellular organisation of carbohydrate metabolism during cold acclimation in different natural accessions of Arabidopsis thaliana. New phytologist 198, 777–787.

Nishizawa A, Yabuta Y, Shigeoka S. 2008. Galactinol and raffinose constitute a novel function to protect plants from oxidative damage. Plant Physiology 147, 1251–1263.

Saito M, Yoshida M. 2011. Expression analysis of the gene family associated with raffinose accumulation in rice seedlings under cold stress. Journal of Plant Physiology 168, 2268–2271.

Schneider T, Keller F. 2009. Raffinose in chloroplasts is synthesized in the cytosol and transported across the chloroplast envelop. Plant Cell Physiology 50, 2174–2182.

Sengupta S, Mukherjee S, Basak P, Majumder AL. 2015. Significance of galactinol and raffinose family oligosaccharide synthesis in plants. Frontiers in Plant Science 6, 656.

Sparkes IA, Runions J, Kearns A, Hawes C. 2006. Rapid transient expression of fluorescent fusion proteins in tobacco plants and genetation of stably transformation plants. Nature Protocols 1, 2019–2025.

Sprenger N, Keller F. 2000. Allocation of raffinose family oligosaccharides to transport and storage pools in Ajuga reptans: the roles of two distinct galactinol synthases. Plant Journal 21, 249–258.

Sui XL, Meng FZ, Wang HY, Wei YX, Li RF, Wang ZY, Hu LP, Wang SH, Zhang ZX. 2012. Molecular cloning, characteristics and low temperature response of raffinose synthase gene in Cucumis sativus L. Journal of Plant Physiology 169, 1883–1891.

Taji T, Ohsumi C, Iuchi S, Seki M, Kasuga M, Kobayashi M, Yamaguchi-Shinozali K, Shinozaki K. 2002. Important roles of drought and cold-inducible genes for galactinol synthase in stress tolerance in *Arabidopsis thaliana*. Plant Journal 29, 417–426.

Tapernoux-L EM, Schneider T, Keller F. 2007. The C-terminal sequence from common bugle leaf galactan:galactan galactosyltransferase is a non-sequence-specific vacuolar sorting determinant. FEBS Letters 581, 1811–1818

Turgeon R, Wolf S. 2009. Phloem transport: cellular pathways and molecular trafficking. Annual Review of Plant Biology 60, 207–221.

Wang CL, Zhang ZP, Miao MM. 2016. SNF1-related protein kinase (SnRK) 1 involved in the regulation of raffinose family oligosaccharide metabolism in cucumber (Cucumis sativus L.) calli. Journal of Plant Growth Regulation 35, 851–864.

Wang DH, Yao W, Song Y, Liu WC, Wang ZZ. 2012. Molecular characterization and expression of three galactinol syntase genes that confer stress tolerance in Salvia miltiorrhiza. Journal of Plant Physiology 169, 1838–1848.

Zhang ZP, Deng YK, Song XX, Miao MM. 2015. Trehalose-6-phosphate and SNF1-related protein kinase 1 are involved in the first-fruit inhibition of cucumber. Journal of Plant Physiology 177, 110–120.

Zuther E, Buchel K, Hundertmark M, Stitt M, Hincha DK, Heyer AG. 2004. The role of raffi nose in the cold acclimation response of *Arabidopsis thaliana*. FEBS Letters 576, 169–173.

